# RNA virus-mediated gene editing for tomato trait breeding

**DOI:** 10.1101/2023.10.01.560115

**Authors:** Mireia Uranga, Verónica Aragonés, Arcadio García, Sophie Mirabel, Silvia Gianoglio, Silvia Presa, Antonio Granell, Fabio Pasin, José-Antonio Daròs

**Affiliations:** Instituto de Biología Molecular y Celular de Plantas (IBMCP), Consejo Superior de Investigaciones Científicas – Universitat Politècnica de València (CSIC-UPV), Valencia 46011, Spain

**Keywords:** Tomato, Potato virus X, CRISPR-Cas9, virus-induced genome editing (VIGE), *STAYGREEN 1* (*SGR1*), *green-flesh* mutant

## Abstract

Virus-induced genome editing (VIGE) is a flexible and robust technology that relies on viral vectors for the transient delivery of CRISPR-Cas components into plants. Tomato (*Solanum lycopersicum* L.) is a mayor horticultural crop grown worldwide; despite its economic importance, little is known about VIGE applicability in this species.

This study presents the successful use of VIGE in tomato for fruit color breeding. We report (i) the generation of a transgenic Cas9-expressing line of tomato cv. Micro-Tom (MT-Cas9), (ii) the use of pLX-PVX, an enhanced RNA viral vector, for single-guide RNA (sgRNA) delivery into MT-Cas9 plants, (iii) heritable, proof-of-concept VIGE of *PHYTOENE DESATURASE* and recovery of albino progeny, and (iv) the recovery of progeny with recolored *green-flesh* fruits by VIGE of *STAYGREEN 1*, thus confirming the successful breeding of tomato fruit color.

Altogether, our results indicate that the presented VIGE approach can be readily applied for accelerated functional genomics of tomato variation, as well as for precision breeding of tomato traits with horticultural interest.

**HIGHLIGHTS:**

- Generation of a transgenic Cas9-expressing line of tomato cv. Micro-Tom
- Use of PVX for sgRNA delivery into Micro-Tom Cas9 plants
- Heritable proof-of-concept VIGE of tomato *PHYTOENE DESATURASE* (PDS)
- Recovery of green-flesh fruits by VIGE of tomato STAYGREEN 1 (SGR1)

## MAIN TEXT

Tomato (*Solanum lycopersicum* L.) is a mayor horticultural crop grown worldwide (https://www.fao.org/faostat/), and a model plant for studying the genetics of fleshy fruit and domestication traits. Modern cultivated tomatoes nonetheless have a narrow genetic basis, which poses a major bottleneck in breeding of new varieties with improved traits^1^.

Owing to their specificity and versatility, clustered regularly interspaced short palindromic repeats (CRISPR)-CRISPR-associated protein (Cas) systems are widely used for efficient genome editing in numerous plant species, including tomato^2^. Current methods in tomato genome editing mainly use the *Streptococcus pyogenes* Cas9 nuclease and single-guide RNA (sgRNA) molecules that are delivered into plant cells through stable *Agrobacterium*-mediated transformation of constructs for the simultaneous Cas9 and sgRNA expression^2^. Stable transformation of CRISPR/Cas9 constructs for multiplexed gene editing allows rapid breeding of tomato traits. Although powerful, this strategy may require careful sgRNA design and combinatorial T-DNA construct assembly, as well as recovery of transgenic plants and complex selection schemes to eventually obtain the desired mutant lines^3^.

The so-called virus-induced genome editing (VIGE) approach uses viral vectors to transiently deliver CRISPR-Cas components into plants. Compared to alternative methods, VIGE (i) allows efficient and fast genome editing for rapid prototyping of sgRNA designs, (ii) minimizes the plant genome integration of exogenous DNA if based on RNA viral vectors, (iii) allows recovery virus-free edited progeny under optimized conditions. VIGE has been successfully applied to the model plants *Nicotiana benthamiana*^4–7^ and *Arabidopsis thaliana*^8^, but not-yet reported for horticultural trait breeding of tomato. Given the VIGE great potential for advanced breeding strategies of this important crop, we developed and describe here a VIGE approach based on an RNA viral vector for tomato genome engineering. We show that the obtained modifications at target genomic loci are heritable, and our VIGE approach can be used for horticultural trait breeding of tomato.

### Proof-of-concept VIGE of tomato *PHYTOENE DESATURASE* (PDS)

We previously reported efficient multiplex, heritable editing in *N. benthamiana* by infection of a line constitutively expressing Cas9 with a potato virus X (PVX; genus *Potexvirus*) vector for sgRNA expression in plant cells^6^. PVX infects a wide range of Solanaceae species. To translate our reported VIGE approach to tomato, we first generated a Cas9-expressing transgenic line of the cultivar Micro-Tom (MT-Cas9) by *Agrobacterium*-mediated stable transformation. T_0_ plants were selected on kanamycin-supplemented medium and by visual tracking of the expression of a reporter gene (DsRed); transgene integration was confirmed by PCR amplification of a T-DNA fragment from plant genomic DNA (Supporting Methods of Supplementary Data). We next engineered pLX-PVX, an enhanced PVX vector based on a mini binary vector of the pLX series^9^ for autonomous replication in *Escherichia coli* and *Agrobacterium*. pLX-PVX allows *Agrobacterium*-mediated inoculation (agroinoculation) of PVX and virus-mediated sgRNA expression in plants (Figure 1A).

**Figure 1.**
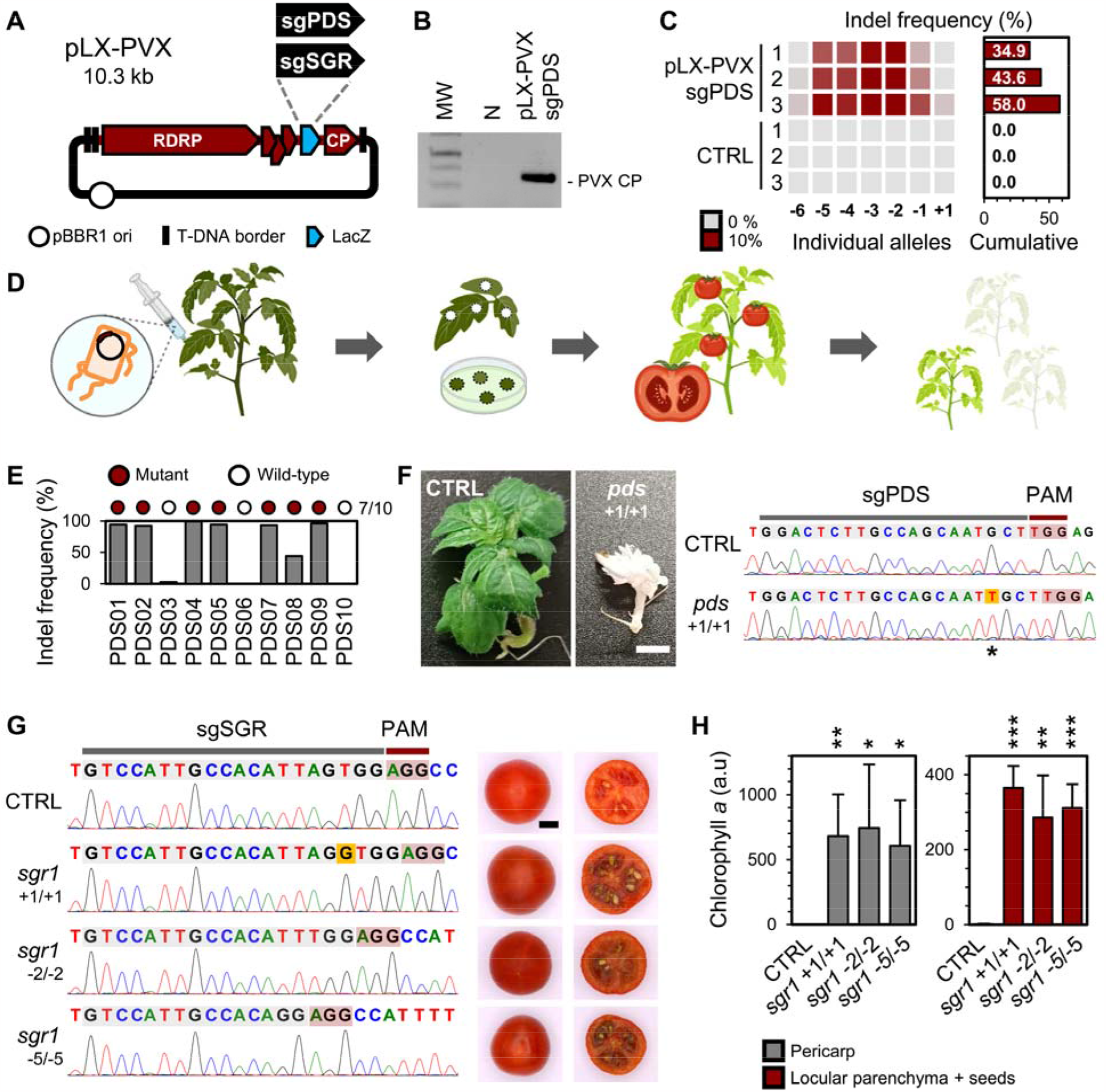
RNA virus/CRISPR-Cas9 editing in tomato. (**A**) Scheme of pLX-PVX, a mini T-DNA vector for agroinoculation of recombinant potato virus X (PVX) constructs for sgRNA delivery. The derivatives pLX-PVX::sgPDS and pLX-PVX::sgSGR include single-guide RNAs (sgRNAs) targeting the tomato *PHYTOENE DESATURASE* (*PDS*; Solyc03g123760) and *STAYGREEN 1* (*SGR1*; Solyc08g080090), respectively. (**B**) *Agrobacterium*-mediated inoculation of pLX-PVX in tomato. RT–PCR detection of a PVX genomic fragment (CP) in a Cas9-expressing tomato cv. Micro-Tom (MT-Cas9) plant agroinoculated with pLX-PVX::sgPDS; samples were collected from upper uninoculated leaves; MW, DNA size standards; N, negative control. (**C**) Virus-induced genome editing (VIGE) of *PDS* in tomato somatic cells. MT-Cas9 plants were agroinoculated with pLX-PVX::sgPDS; after 6 weeks, VIGE was assessed in upper uninoculated leaf samples by PCR of a *PDS* genomic fragment and Sanger trace deconvolution. Indel frequency percentages are shown; CTRL, uninoculated MT-Cas9 plants. (**D**) Experimental workflow for tomato trait breeding using the PVX/CRISPR-Cas9 system. sgRNAs are delivered into MT-Cas9 by pLX-PVX agroinoculation, whole plants are regenerated *in vitro* from upper uninoculated leaf tissue, and progeny screened for mutant line identification. (**E**) VIGE of *PDS* in regenerated tomato plants. Indel frequency percentages and genotypic classification of 10 regenerated plants are shown. (**F**) Rescue of homozygous *PDS* mutant progeny. Phenotype and Sanger trace of the progeny line PDS02.3 with a homozygous loss-of-function *PDS* allele (*pds* +1/+1); CTRL, control condition. (**G**) Rescue of homozygous *SGR1* mutant progeny. Sanger traces and mature fruit phenotypes of progeny lines with homozygous loss-of-function *SGR1* alleles; CTRL, unedited MT-Cas9. (**H**) Chlorophyll quantification in fruits of *SGR1* mutants. Normalized fluorometric amounts of chlorophyll *a* in mature fruit samples are plotted (mean ± SD, *n* = 4); significance levels versus the control unedited MT-Cas9 (CTRL) as per Student’s *t*-test are shown; *, *p* _≤_ 0.05; **, *p* _≤_ 0.01; ***, *p* _≤_ 0.001.

We reasoned that PVX-mediated sgRNA expression in MT-Cas9 would result in editing events of the tomato genome. For proof-of-concept VIGE assays, we assembled pLX-PVX::sgPDS for expression of a sgRNA targeting *PHYTOENE DESATURASE* (*PDS*; Solyc03g123760) (Figure S1A; Table S1), whose loss-of-function mutants have a reported photobleaching phenotype^2^. Cotyledons of MT-Cas9 seedlings were agroinoculated with pLX-PVX::sgPDS, and samples were collected from upper uninoculated leaves after 6 weeks. PVX infectivity in inoculated plants was confirmed by RT–PCR detection of a viral genomic fragment (Figure 1B). Although no photobleaching could be observed (Figure S1B), PCR amplification of a genomic fragment spanning the sgRNA target site followed by Sanger trace deconvolution revealed the presence of multiple *PDS* alleles with insertion-deletion (indel) mutations and cumulative indel frequencies of 34.9-58.0% (Figure 1C; Supporting Methods). These results confirm that RNA virus-mediated sgRNA delivery can produce somatic mutations in a tomato Cas9-expressing line.

Given the high editing efficiency detected in somatic cells, we conceived an experimental workflow for heritable trait breeding using the PVX/CRISPR-Cas9 system in tomato (Figure 1D). At 21 dpi, we collected upper uninoculated leaves of infected plants, detected *PDS* indel frequencies of 19.7-46.9% (x□ = 28.5; Figure S1C), and regenerated whole plants by tissue culture (Figure 1D; Supporting Methods). Genotyping of randomly picked green plants indicated that 70% (7/10) of them had *PDS* mutant alleles (Figure 1E). Among these, the line PDS02 had a 1-bp insertion allele (+1) and a 3-bp deletion allele (−3) in *PDS* (Table S2). To confirm the mutation heritability, fruits were collected from the self-pollinated PDS02 line and progeny analyzed. The recovered lines PDS02.1 and PDS02.2 were bi-allelic (+1/-3) or homozygous (−3) mutants showing the green, wildtype phenotype (Figure S1D; Table S2). Remarkably, the progeny line PDS02.3 showed a photobleaching phenotype and the homozygous +1 allele (*pds* +1/+1; Figure 1F; Table S2), which caused the *PDS* complete inactivation. These findings demonstrate that our VIGE approach yields, through tissue culture regeneration, heritable editing events and can be used to recover loss-of-function mutant progeny.

### VIGE of tomato *STAYGREEN 1* (SGR1) yields fruits with the *green-flesh* phenotype

Heirloom accessions show a wide range of variation in fruit shape, size, and color. Among others, Purple Calabash, Cherokee Chocolate, Black Krim, and Black Prince are mutants of *STAYGREEN 1* (*SGR1*; Solyc08g080090)^10^. SGR1 inactivation inhibits degradation of chlorophyll *a*; simultaneous chlorophyll and lycopene accumulation during ripening produces a *green-flesh* phenotype characterized by red-brown fruits^3,10,11^.

Supported by the *PDS* editing results, we wondered if our VIGE approach could be used for *SGR1* editing and breeding of fruit color. We thus assembled pLX-PVX::sgSGR to target *SGR1* (Figures 1A, S2A; Table S1), and agroinoculated it into MT-Cas9 seedlings. At 21 dpi, Sanger trace deconvolution analysis revealed an average *SGR1* indel frequency of 21.4% in upper uninoculated leaves (Figure S2B), which were then used for in-vitro regeneration of whole plants. 60% (9/15) of them had *SGR1* mutant alleles (Figure S2C). Subsequent fruit collection and progeny analysis allowed identifying homozygous lines with a single mutant allele of *SGR1* each (*sgr1* +1/+1, *sgr1* −2/−2, and *sgr1* −5/−5), and whose fruits displayed the *green-flesh* phenotype (Figure 1G); no PVX could be detected in these lines (Figure S3). Fluorometric and absorbance-based quantification revealed a significant increase of chlorophyll *a* amount in the pericarp and locular parenchyma plus seed samples of *sgr1* mutant lines compared to the control, unedited MT-Cas9 (Figures 1H, S4A). Results of two complementary methods were highly consistent (*r* = 0.967, *p* < 0.0001; Figure S4B), overall confirming the SGR1 inactivation. Altogether, our VIGE results prove the successful recovery of virus-free, *SGR1* loss-of-function mutants and thus horticultural trait breeding of tomato.

### Concluding remarks

In conclusion, we reported here (i) generation of a transgenic Cas9-expressing line of tomato cv. Micro-Tom (MT-Cas9), (ii) use of pLX-PVX, an enhanced RNA viral vector, for sgRNA delivery into MT-Cas9 plants, (iii) heritable, proof-of-concept VIGE of *PHYTOENE DESATURASE* and recovery of albino progeny, and (iv) recovery of *green-flesh* fruits by VIGE of *STAYGREEN 1*, altogether, indicating that the presented VIGE approach can be readily applied for breeding of tomato traits with horticultural interest.

Zhenghe Li and co-workers recently reported a tospovirus vector system for viral delivery of Cas nucleases and their corresponding CRISPR guide RNAs, which enabled VIGE in a variety of non-transgenic crops^12^. The system may however be unsuitable for modern tomato varieties with *Sw-5*, granting broad-spectrum resistance against tospoviruses^13^, which could be nonetheless amenable for the PVX-based VIGE approach here presented. In sum, our work reinforces VIGE as a flexible and robust technology for accelerating functional genomics of tomato variation^1^, as well as for precision breeding of novel traits to tackle global food supply and health problems^14,15^.

## ACKNOWLEDGEMENTS

This work was supported by grant PID2020-114691RB-I00 from Ministerio de Ciencia e Innovación (Spain) through the Agencia Estatal de Investigación. An.G. received funding form Horizon 2020 with HARNESSTOM contract 101000716. F.P. is supported by the “Juan de la Cierva Incorporación” contract IJC2019-039970-I from Ministerio de Ciencia e Innovación (Spain), S.G. by CIAPOS/2021/316 from Generalitat Valenciana (Spain) and European Social Fund, and Ar.G. by the fellowship FPU20/05477 from Ministerio de Ciencia e Innovación (Spain).

## AUTHOR CONTRIBUTIONS

All authors participated in the work conception and design. M.U., V.A., A.G., S.M., S.G. and S.P. performed the experiments. All authors participated in the result and data analyses. M.U., F.P. and J.-A.D. wrote the manuscript with input from the rest of the authors.

## DATA AVAILABILITY STATEMENT

Supplementary data accompany this article. The pLX-PVX vector is available at Addgene with catalog number (*pending*). The sequence information of tomato *PDS* (Solyc03g123760) and *SGR1* (Solyc08g080090) are available at Solanaceae Genomics Network (https://solgenomics.net/). Further information and requests for resources should be directed to and will be fulfilled by the corresponding authors.

## CONFLICT OF INTEREST

The authors declare no conflict of interest.

## SUPPLEMENTARY DATA

Supplementary data include: Supporting methods, Supplementary references, Tables S1-S3, and Figures S1-S4.

## Notes

### Competing Interest Statement

The authors have declared no competing interest.

