## SUPPLEMENTARY DATA for "RNA virus-mediated gene editing for tomato trait breeding"

### TABLE OF CONTENTS

**SUPPORTING METHODS**

**SUPPLEMENTARY REFERENCES**

**SUPPLEMENTARY TABLES**

- **Table S1. Single-guide RNAs used in the study**
- **Table S2. Heritability of *PHYTOENE DESATURASE* (*PDS*) mutations**
- **Table S3. Primers used in the study**

**SUPPLEMENTARY FIGURES**

- **Figure S1. Virus-induced genome editing (VIGE) of tomato *PHYTOENE DESATURASE* (*PDS*)**
- **Figure S2. VIGE of tomato *STAYGREEN 1* (*SGR1*)**
- **Figure S3. Chlorophyll spectrophotometric quantification in fruits of *SGR1* mutant lines**
- **Figure S4. PVX detection in *SGR1* mutant lines**

### SUPPORTING METHODS

#### Plasmid construction

The binary vector GB2234 for plant expression of a codon-optimized *Streptococcus pyogenes* Cas9 (hCas9) sequence under the control of the cauliflower mosaic virus (CaMV) 35S promoter and *Agrobacterium tumefaciens* nopaline synthase terminator ($P_{35S}$:hCas9:$T_{nos}$) was reported[1]. GB2234 includes a kanamycin resistance gene (nptII) and a red fluorescent protein gene (DsRed) for plant selection and its sequence is available at the GoldenBraid database (https://gbcloning.upv.es/feature/GB_UA_2578/)[2].

pLX-PVX was used for recombinant viral vector assembly. Potato virus X (PVX) genomic cDNA were amplified from pPVX[1], a pgR107-derivative based on pGreen0000 which lacks autonomous replication in *Agrobacterium* cells[3,4]. The obtained viral sequence was inserted into pLX-B2 (GenBank: KY825137; Addgene: 160636), a mini T-DNA binary vector of the pLX series[5] for autonomous replication in *Escherichia coli* and *Agrobacterium*. pLX-PVX T-DNA includes a full-length potato virus X (PVX) cDNA flanked by the CaMV 35S promoter and *A. tumefaciens* nos terminator, sgRNA expression is driven by the PVX coat protein (CP) promoter, whereas PVX CP is expressed from a heterologous promoter derived from bamboo mosaic virus. pLX-PVX includes the LacZ reporter for white-blue screen of recombinant vectors.

The single-guide RNA (sgRNA) sequences sgPDS and sgSGR used to respectively target the tomato *PHYTOENE DESATURASE* (PDS; Solyc03g123760.2.1) and *STAYGREEN 1* (SGR1; Solyc08g080090.2) are shown in Table S1. Recombinant pLX-PVX derivatives for sgRNA expression were assembled using standard molecular biology techniques[6]. Briefly, fragments including sgPDS and sgSGR fused to a truncated *Arabidopsis thaliana FLOWERING LOCUS T* sequence[7] were obtained by PCR reactions including high-fidelity Phusion DNA polymerase (Thermo Fisher Scientific), pPVX::sgXT2B-tFT as the DNA template[1], and the primer pairs D4747/D3898 and D4748/D3898, respectively (Table S3). Gel purified fragments were inserted into MluI-linearized pLX-PVX by Gibson assembly reactions with the NEBuilder HiFi DNA assembly master mix (New England Biolabs). Recombinant plasmids were purified from white *E. coli* colonies selected on agar medium plates supplemented with kanamycin and 5-Bromo-4-chloro-3-indolyl β-D-galactopyranoside (X-Gal). The obtained viral vectors pLX-PVX::sgPDS and pLX-PVX::sgSGR were verified by Sanger sequencing.

#### Plant materials

Stable transformation of the tomato cultivar 'Micro-Tom'[8] was done using the binary vector GB2234 as described[9]. Micro-Tom transformants (MT-Cas9) were identified by kanamycin selection and monitoring of the DsRed reporter expression under an epifluorescence stereoscope (Leica DMS1000). Genomic DNA was purified from plant samples using silica gel columns, and stable transgene integration was confirmed by PCR amplification of a 792-bp T-DNA fragment using the primer pair D3663/D3664 (Table S3) and. MT-Cas9 plants were grown in greenhouse chambers at ~24°C under a 16-h-day/8-h-night photoperiod.

**Viral vector agroinoculation and analysis of virus infectivity**

*A. tumefaciens* C58C1 cells including a disarmed pTi were electroporated with viral vector plasmids; colonies were selected on plates supplemented with rifampicin and kanamycin. Suspensions of the transformed bacteria were prepared as described[6] and used to inoculate fully expanded cotyledons from 10-day-old tomato plants.

Total RNA was purified from upper non-inoculated leaf samples using silica gel columns (Zymo Research) as previously described[6]. cDNA was synthesized using RevertAid reverse transcriptase (Thermo Fisher Scientific) and the primer D2409 (Table S3); the PVX coat protein (CP) sequence was amplified in PCR reactions including the obtained cDNA, Thermus thermophilus DNA polymerase (Biotools) and the primer pair D2410/D3436 (Table S3). RT-PCR products were analyzed by electrophoresis in 1% agarose gels in TAE buffer (40 mM Tris, 20 mM sodium acetate, and 1 mM EDTA, pH 7.2), and visualized by ethidium bromide staining.

***In vitro* regeneration of tomato plants**

At 21 days post inoculation (dpi), the first and second systemic leaves showing symptoms of viral infection were detached, surface-sterilized by submersion in sterilization solution (10% bleach, 0.02% Nonidet P-40) for 10 min, and then rinsed three times in sterile water. One-cm leaf slices were obtained and plated on to callus induction medium including 1× Murashige & Skoog salts (MS) with vitamins, 0.5 g/L 2-(N-Morpholino)ethanesulfonic acid hydrate (MES), 20 g/L glucose, 10 g/L phytoagar, 0.1 mg/L 3-indolyl acetic acid (IAA), 0.75 mg/L trans-zeatin, pH 5.7. Leaf slices were transferred to fresh plates every 2-3 weeks until shoots emerged. The shoots were then transferred to elongation medium (1× MS with vitamins, 0.5 g/L MES, 20 g/L glucose, 10 g/L phytoagar, 0.1 mg/L trans-zeatin, pH 5.7) for 1-2 weeks until they reach a length of 1-2 cm. Subsequently, shoots were cut and transferred to rooting medium (0.5× MS with vitamins, 0.5 g/L MES, 20 g/L glucose, 10 g/L phytoagar, 0.1 mg/L IAA, pH 5.7). Once rooted plantlets reached a height of >5 cm, they were transferred to soil and covered with plastic cups to keep moisture in. After acclimation to soil conditions, plastic cups were removed for adequate plant growth. Following flowering and fruit development, fruits were harvested from each plant for progeny analysis.

**Genome editing analysis**

Genomic DNA was purified from leaf samples using silica gel columns[6]. PCR using high-fidelity Phusion DNA polymerase (Thermo Fisher Scientific), and the primer pairs D4762/D4763 and D4766/D4767 (Table S3) were respectively used to amplify genomic fragments spanning the sgPDS and sgSGR target sites. PCR products were separated by agarose gel electrophoresis, purified, and subjected to Sanger sequencing. Genome editing efficiency was quantified by Sanger trace deconvolution[6] using TIDE (http://shinyapps.datacurators.nl/tide-batch/)[10] or ICE (https://ice.synthego.com/)[11].

**Chlorophyll quantification**

Samples were collected from red mature tomato fruits. Aliquots of pericarp tissue (±100 mg) and seeds plus the surrounding locular parenchyma (±50 mg) were respectively mixed with 10 or 20 volumes (~1 mL) of aqueous 80% (v/v) acetone precooled to –20°C, homogenized using a ball mill (Star-Beater, VWR) for 5 min at 30 $s^{-1}$, and mixed (1 h) at 250 rpm at 4°C in the dark. Samples were centrifuged at 14,000×*g* (5 min) to remove cell debris; supernatants were collected and kept in the dark until analysis.

Fluorometric quantification of chlorophyll *a* (chl *a*) was done in a monochromator-based plate reader (Infinite M Plex, Tecan Group) as reported[12], with minor modifications . Briefly, the above-prepared supernatant samples were mixed 1:1 with aqueous 80% (v/v) acetone or aqueous 80% (v/v) acetone, 0.02 N HCl, to obtain native (*N*) and acidified (*A*) extracts, respectively. Samples were aliquoted to a black 96-well flat-bottom plate (Nunc), and top reading measurements recorded using $\lambda_{Ex}$ 436 nm and $\lambda_{Em}$ 680 nm; the instrument gain value was manually adjusted to avoid signal saturation. Fluorescence intensity of *N* and *A* extracts was measured to obtain $F_N$ and $F_A$, respectively; chl *a* amount was calculated as $F_N - F_A$ with background values subtracted.

Spectrophotometric quantification of chl *a* was done as reported[13]. Briefly, supernatant absorbance was measured at 646 nm ($A_{646}$) and 663 nm ($A_{663}$) against an aqueous 80% (v/v) acetone blank in a plate reader (Infinite M Plex, Tecan Group), and chl *a* quantified using the equation:

Chl *a* = $12.21 \cdot A_{663} - 2.81 \cdot A_{646}$

**Statistics**

Student's *t*-test was used for two-group comparisons; significance levels of *p* values are indicated in the figures. Pearson correlation coefficient was calculated to estimate the linear relationship between the two variables.

### SUPPLEMENTARY TABLES

#### Table S1. Single-guide RNAs used in the study

| sgRNA | Target gene | Target gene accession number | Sequence | Ref. |
|---|---|---|---|---|
| sgPDS | PHYTOENE DESATURASE | Solyc03g123760 | GGACTCTTGCCAGCAATGCT | [14] |
| sgSGR | STAYGREEN 1 | Solyc08g080090 | GTCCATTGCCACATTAGTGG | [9] |

#### Table S2. Heritability of *PHYTOENE DESATURASE* (*PDS*) mutations

| Line[a] | Allele | Allele sequence[b] | Allele frequency[c] |
|---|---|---|---|
| Control | wild-type | GGACTCTTGCCAGCAATGCTTGG | — |
| PDS02 | +1 | GGACTCTTGCCAGCAATTGCTTGG | 47% |
| | -3 | GGACTCTTGCCAGC---GCTTGG | 44% |
| PDS02.1 | +1 | GGACTCTTGCCAGCAATTGCTTGG | 48% |
| | -3 | GGACTCTTGCCAGC---GCTTGG | 46% |
| PDS02.2 | -3 | GGACTCTTGCCAGC---GCTTGG | 74% |
| PDS02.3 | +1 | GGACTCTTGCCAGCAATTGCTTGG | 97% |

[a] The *in vitro* regenerated line PDS02 and its progeny lines PDS02.1, PDS02.2, and PDS02.3 are shown

[b] sgPDS target sequence with indels in mutant alleles shown in red; the protospacer adjacent motif (PAM) is underlined

[c] Frequencies by Sanger trace deconvolution

#### Table S3. Primers used in the study

| Primer | Primer sequence (5'→3') | Purpose |
|---|---|---|
| D4747 | GAGGTCAGCACCAGCTAGCAGGACTCTTGCCAGCAATGCTGTTTTAGAGC | sgRNA cloning |
| D4748 | GAGGTCAGCACCAGCTAGCAGTCCATTGCCACATTAGTGGGTTTTAGAGC | sgRNA cloning |
| D3898 | GGGAAACTTAACAAACCCTATTGGCCATAAGTAACCTTTAG | sgRNA cloning |
| D2409 | ATTTATATTATTCATACAATCAAACC | PVX detection |
| D3436 | ATGTCAGGCCTGTTCACTATCC | PVX detection |
| D2410 | TGGTGGTGGTAGAGTGACAAC | PVX detection |
| D3663 | ATGATTGAACAAGATGGATTGC | Plant genotyping |
| D3664 | GAAGAACTCGTCAAGAAGGCGA | Plant genotyping |
| D4762 | TCTGTATTTTGTCTAGCTTCTCCT | Plant genotyping |
| D4763 | TGCACTGCATTGAAAGTTCGTC | Plant genotyping |
| D4766 | TCCGTGCCGTCTATTGTTCT | Plant genotyping |
| D4767 | ACCCAAACTAAAGCTTCTTGTAACT | Plant genotyping |

### SUPPLEMENTARY FIGURES

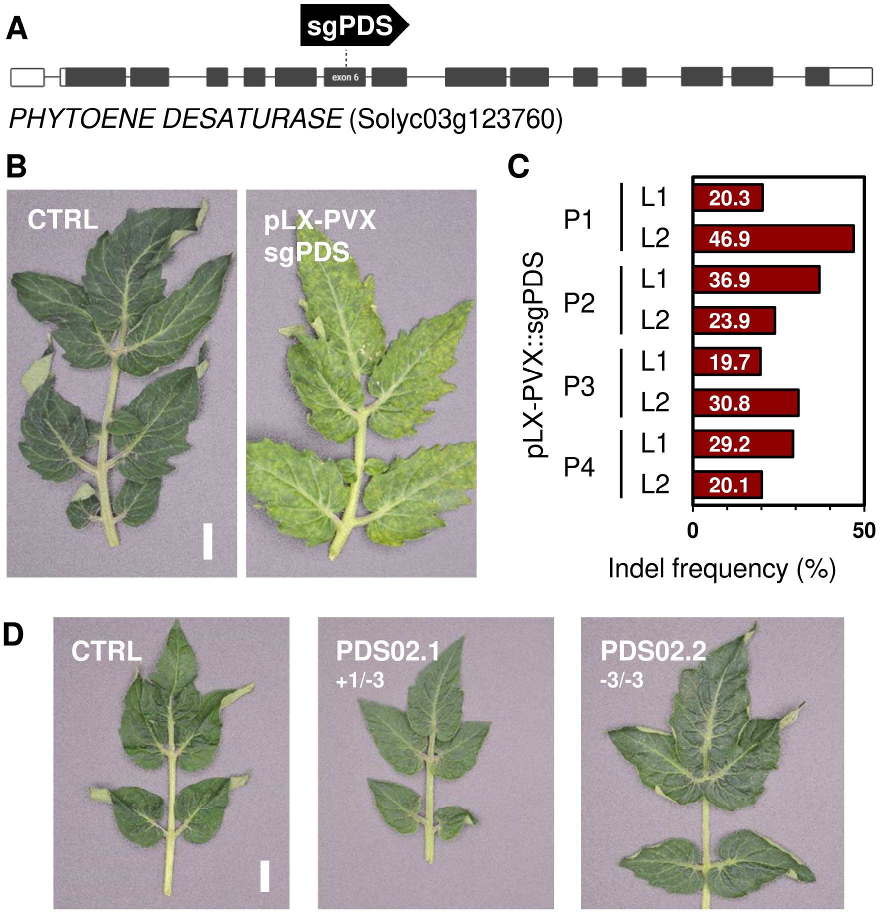


**Figure S1. Virus-induced genome editing (VIGE) of tomato *PHYTOENE DESATURASE* (*PDS*).** (**A**) Genetic structure of the tomato *PDS* (Solyc03g123760) and the pLX-PVX::sgPDS target region in exon 6. (**B**) Phenotype of a Cas9-expressing tomato cv. Micro-Tom (MT-Cas9) plant after 6 weeks of agroinoculation with pLX-PVX::sgPDS; CTRL, untreated plant; scale = 1 cm. (**C**) VIGE of *PDS* in tomato somatic cells. MT-Cas9 plants were agroinoculated with pLX-PVX::sgPDS; after 21 days, VIGE was assessed in upper uninoculated leaf samples by PCR of a *PDS* genomic fragment and Sanger trace deconvolution. Indel frequency percentages are shown for four plants (P1 – P4) and two leaves each (L1 – L2). (**D**) Pictures show the phenotypes of PDS02.1 and PDS02.2, progeny lines of a single plant (PDS02) regenerated from MT-Cas9 agroinoculated with pLX-PVX::sgPDS; see Table S2 for genotyping results.

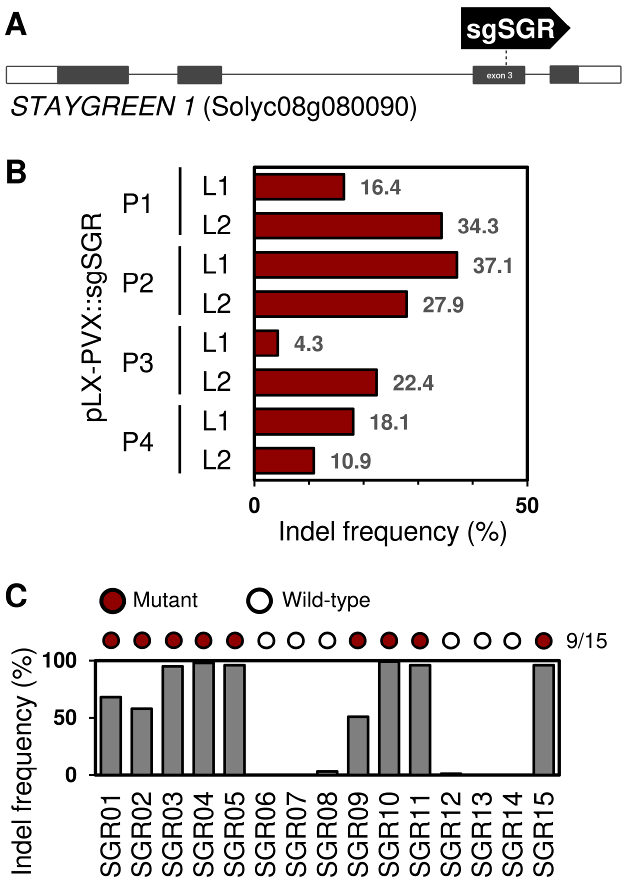


**Figure S2. VIGE of tomato *STAYGREEN 1* (*SGR1*).** (**A**) Genetic structure of the tomato SGR1 (Solyc08g080090) and the pLX-PVX::sgSGR target region in exon 3. (**B**) VIGE of *SGR1* in tomato somatic cells. MT-Cas9 plants were agroinoculated with pLX-PVX::sgSGR; after 21 days, VIGE was assessed in upper uninoculated leaf samples by PCR of an *SGR1* genomic fragment and Sanger trace deconvolution. Indel frequency percentages are shown for four plants (P1 – P4) and two leaves each (L1 – L2). (**C**) VIGE of *SGR1* in regenerated tomato plants. Indel frequency percentages and genotypic classification of 15 regenerated plants are shown.

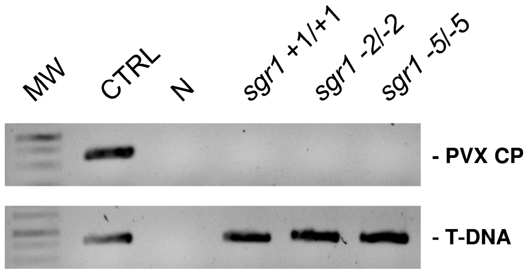


**Figure S3. PVX detection in *SGR1* mutant lines.** Top, RT–PCR detection of a PVX genomic fragment (CP) from total RNA samples; bottom, PCR detection of a T-DNA fragment from genomic DNA samples; MW, DNA size standards; CTRL, positive control; N, negative control.

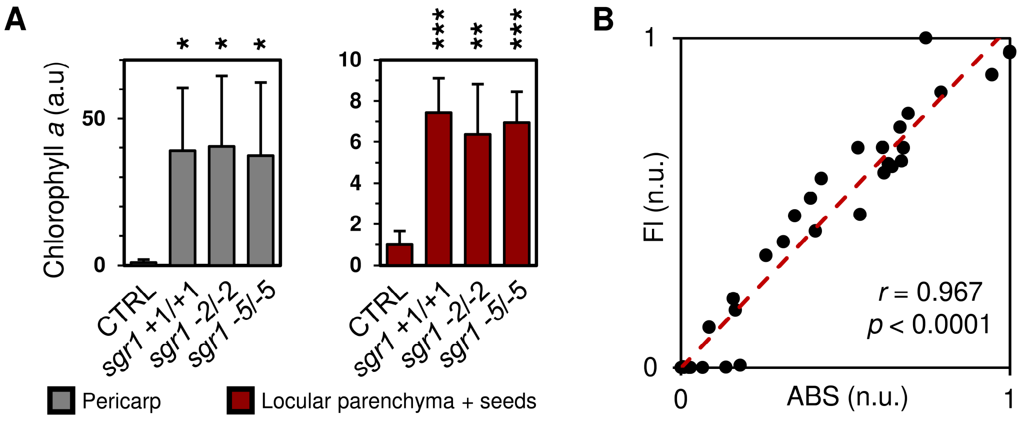


**Figure S4. Chlorophyll spectrophotometric quantification in fruits of *SGR1* mutant lines.** (**A**) Chlorophyll spectrophotometric quantification. Normalized absorbance-derived amounts of chlorophyll *a* in fruit samples are plotted (mean ± SD, $n = 4$); samples from the unedited MT-Cas9 line were used as the control condition (CTRL); significance levels versus CTRL as per Student's *t*-test are shown; *, $p \leq 0.05$; **, $p \leq 0.01$; ***, $p \leq 0.001$. (**B**) Linear relationship between chlorophyll *a* normalized units (n.u.) determined by spectrophotometric (ABS) and fluorometric (FI) analyses ($n = 32$); Person's *r* and *p* value are indicated.
